# Cell-intrinsic and -extrinsic functions of the ESCRT-III component Shrub in cytokinetic abscission of *Drosophila* Sensory Organ precursor

**DOI:** 10.1101/2022.10.27.514017

**Authors:** Céline Bruelle, Mathieu Pinot, Emeline Daniel, Marion Daudé, Juliette Mathieu, Roland Le Borgne

## Abstract

While the molecular mechanisms underlying the abscission of isolated cells are largely decrypted, those of fast-cycling, epithelial progenitors surrounded by epidermal cells (ECs) connected by junctions remain largely unexplored. Here, we investigated the remodeling of the permeability barrier ensured by septate junctions (SJs) during cytokinesis of Drosophila sensory organ precursor (SOP). We report that SOP cytokinesis involves the coordinated polarized assembly and remodeling of SJs in the dividing cell and its neighbors, which remained connected via membrane protrusions pointing toward the SOP midbody. SJs assembly and midbody basal displacement occur more rapidly in SOP than in ECs, leading to a faster disentanglement of the protrusions that precedes midbody release. As reported in isolated cells, the endosomal sorting complex required for transport-III component Shrub/CHMP4B is recruited at the midbody and cell-autonomously regulates abscission. In addition, we found that Shrub is recruited to membrane protrusions, is required for SJ integrity, and that alteration of SJ integrity leads to premature abscission. Our study uncovers cell-intrinsic and -extrinsic functions of Shrub in epithelial abscission to support the coordination of permeability barrier maintenance and abscission in SOPs.

## Introduction

Cell junctions are essential for the chemical and mechanical functions of epithelia (Higashi et al., 2016). Adherens junctions (AJs) and Tight junctions (TJ) in vertebrates/ Septate Junctions (SJs) in invertebrate ensure the mechanical and permeability barriers respectively (Banerjee et al., 2006; Harris and Tepass, 2010; Shin et al., 2006; Tepass et al., 2001; Tsukita et al., 2001). *Drosophila* bicellular SJs (bSJs) are composed of a highly stable core complex containing more than 20 proteins including cytosolic proteins such as Coracle (Cora) and Disc-large (Dlg), Claudin-like proteins including Kune Kune, cell adhesion molecules such as Fasciclin III (Fas 3) transporters such as Na+/K+ ATPase alpha (ATPα) and beta (Nervana2 (Nrv2)) subunits (Faivre-Sarrailh, 2020; Genova and Fehon, 2003; Izumi and Furuse, 2014; Kaplan, 2002; Nelson et al., 2010; Snow et al., 1989; Ward et al., 1998). The localization of SJ components are interdependent of each other (Oshima and Fehon, 2011).

While cells need to establish and maintain proper permeability barrier, SJs remain highly plastic, especially during cell division, a fundamental process for the development and function of all organs. Abscission is the final step of cytokinesis that leads to the physical separation of the two daughter cells and has been proposed to help regulate cell fate acquisition (Chaigne et al., 2020; Ettinger et al., 2011). Cytokinesis is initiated at the onset of anaphase and begins with the formation of an actomyosin ring that constricts to ultimately form the midbody (Fededa and Gerlich, 2012; Glotzer, 2005; Green et al., 2012). The midbody present at the intercellular bridge connects the two daughter cells, and recruits effectors of abscission. In *Drosophila*, the centralspindlin protein complex recruits Alix to the midbody which in turn recruits the endosomal sorting complex required for transport-III (ESCRT III) component Shrub (the CHMP4B ortholog). Shrub/CHMP4B have the property to form homopolymers that are proposed to drive abscission (Eikenes et al., 2015; Guizetti et al., 2011; Lie-Jensen et al., 2019; Matias et al., 2015). In addition to being required for multivesicular body formation and abscission, CHMP4B exerts different functions including viral budding and, plasma and nuclear membrane resealing (Agromayor and Martin-Serrano, 2013; Lie-Jensen et al., 2019; Vietri et al., 2020). While there is a great deal of knowledge on the molecular mechanisms underlying abscission of isolated cells, abscission in tightly packed polarized epithelial cells possessing junctional complexes remains largely under-explored.

Several studies have described that the formation of the new AJs is coordinated with the early steps of cytokinesis leading to the positioning of the midbody basal to the newly formed adhesive interface (Firmino et al., 2016; Founounou et al., 2013; Guillot and Lecuit, 2013; Herszterg et al., 2013; Higashi et al., 2016; Lau et al., 2015; Morais-de-Sá and Sunkel, 2013). It has also been showed that the neighbor cells maintain SJ contact with the dividing cell through SJ finger-like protrusions connected to the midbody in the *Drosophila* pupal notum and wing imaginal disc (Daniel et al., 2018; Wang et al., 2018). Novel SJ assembles below the AJs and spread basally concomitantly with the basal displacement of the midbody. Once the new SJ is formed, the midbody leaves the SJ about 1.5 hours after the onset of anaphase and abscission is taking place more than 5 hours post anaphase (Daniel et al., 2018; Wang et al., 2018). The sequence of events leading to abscission and junction remodeling is proposed to ensure the maintenance of the epithelial barrier functions in proliferative epithelia. This issue remains poorly characterized in epithelia composed of cells with distinct identities.

*Drosophila* notum consists of a single-layer epithelium with two types of cells, epidermal (EC) and sensory organs (SO) (Fig. 1A). Sensory organs precursor (SOP) divides asymmetrically to generate pIIa and pIIb cells that undergo subsequent cell divisions giving rise to the four cells composing the adult SO (Fig. 1A’) (Fichelson and Gho, 2003; Gho et al., 1999; Hartenstein and Posakony, 1989). At each cell division, cell fate determinant Numb and Neuralized are unequally segregated to control Notch-dependent binary fate acquisition (Langevin et al., 2005; Le Borgne and Schweisguth, 2003; Rhyu et al., 1994). In contrast to EC, SO and daughters are fast cycling cells (Audibert, 2005) (Fig. 1A’) raising the question of how permeability barrier and abscission occur during SOP cytokinesis to ensure the proper progression of cell divisions while enabling cell fate determination. In this study, we first characterized the coordination between the midbody basal displacement and the SJ remodeling during SOP cytokinesis. We next investigated the functions of Shrub and SJ components in abscission. We provided evidence that SJ components negatively regulate the timing of abscission during SOP cytokinesis. We also report that Shrub, by controlling the abscission and the remodeling of SJs, exerts two complementary functions to orchestrate SO abscission.

**Fig. 1.**
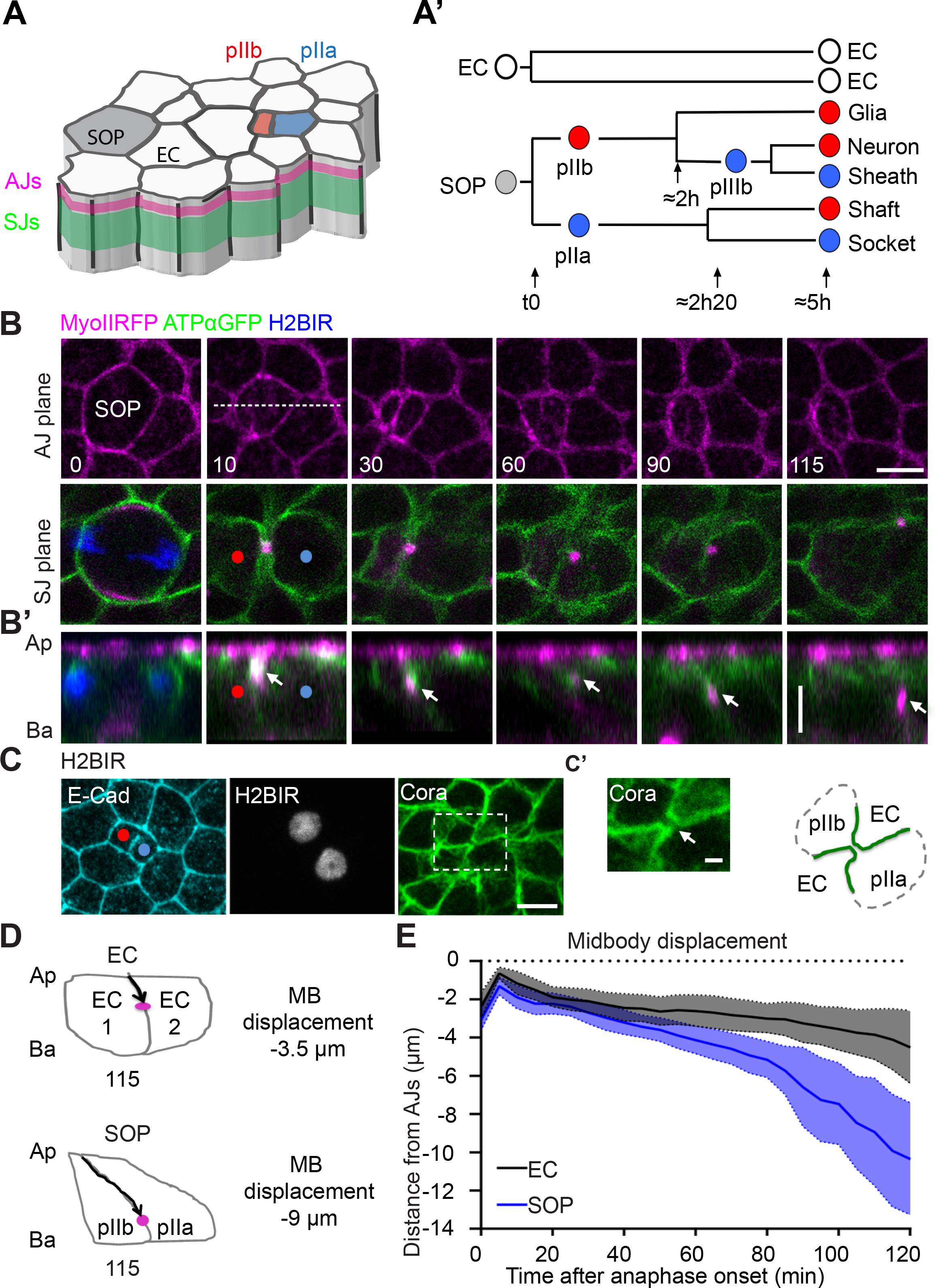
Midbody assembly and basal displacement throughout epithelial cell cytokinesis. (A) Schematic representation of the *Drosophila* pupal notum composed of epidermal cells (white) and sensory organ precursors (SOP, grey). Adherens junctions (AJ) and septate junctions (SJ) were represented in magenta and green respectively. (A’) EC daughter cells cytoplasmic isolation happens after 5 hours after anaphase onset of their mother. In the meantime, SOP undergo four rounds of asymmetric cell division. 2h and 2h20 refers to the approximative time at which the respective daughter cells start to divide. At each division, fate identity relies on differential activation of Notch (Notch is activated in the cells depicted in blue). (B) Time-lapse of SOP cells (n=12) expressing MyoIIRFP (magenta), ATPαGFP (green) and Histone 2B IR (H2BIR) to identify the SOP (blue). B’ correspond to the orthogonal views along the white dashed lines depicted at t=10 min on panel B. White arrows point to the midbody. AJ, adherens junction; SJ, septate junction; Ap: Apical; Ba: Basal. (C) Localization of E-Cad (cyan) and Cora (green) at the pIIa-pIIb cell stage identified using H2BIR (white). Higher magnifications in (C’) correspond to the ROI in the white dotted line square depicted in (C) and white arrows show the finger-like protrusions. (n>3, 3 nota per condition). Right: Schematic representation of the SOP with the finger like protrusions in green as in (C’). (D) Schematic representation of the midbody basal displacement in EC and SOP corresponding to t115 (1A’ and 1B’). MB: Midbody (E) Plot of the quantification of the apical to basal midbody displacement over time relative to AJs in EC (black, n=40) and SOP (blue, n=30) dividing cells. ANCOVA, p-value < 2.2.10^−16^ ***. Solid lines represent simple linear fit. Time is in minute with t= 0 corresponding to anaphase onset (B-B’, E). Scale bars represent 5 μm (B-C) and 1 μm (C’). Red and blue dots mark the pIIb and pIIa respectively (B-C).

## Results

### Basal displacement of the midbody is faster in SOP than in epidermal cells

To monitor SOP cell cytokinesis and compare it to that of epidermal cells (ECs), we live-imaged non-muscle Myosin II light chain tagged with RFP (MyoIIRFP) to label the plane of AJs and the actomyosin contractile ring dynamics, ATPα tagged with GFP (ATPαGFP) to monitor the SJs (Fig. 1B-B’ and Fig. S1A-A’). SOP and its pIIa/pIIb daughter cells were identified using Histone 2B∷IRFP670 (H2BIR) expressed under the minimal promoter of *neuralized* (Fig. 1B-1B’). In each movie, time is expressed in min, with t0 corresponding to the onset of anaphase. In both EC and SOP, the constriction of the actomyosin contractile ring gives rise to the midbody located basal to the AJ, within the SJs (Fig. 1B-B’, Fig.S1A-A’ at t10 and (Daniel et al., 2018; Founounou et al., 2013; Guillot and Lecuit, 2013; Herszterg et al., 2013; Morais-de-Sá and Sunkel, 2013)). As described for dividing ECs (Fig. 1-Fig. S1A, t10 ;(Daniel et al., 2018; Wang et al., 2018)), as a result of membrane ingression induced by actomyosin ring pulling, neighboring cells remain tightly associated to SOPs by finger-like protrusions, marked with the SJs components ATPαGFP and Cora (Fig. 1B, t10; SJ plane, Fig. 1C-C’, t10; SJ plane). While another commonality of the SOPs and ECs is the displacement of the midbody towards the basal pole (Fig. 1B’, Fig. S 1A’, arrows), we found that the basal displacement of the SOP midbody is 1.75 times faster (2.8±0.1 μm/h) than that of the EC (1.6±0.1 μm/h) in the 80 min after anaphase onset (Fig. 1D-E). Then, we observed an inflection in the curve, the SOP midbody being displaced with a faster kinetic towards the basal side (6.9±0.2 μm/h) (Fig. 1E), as it is progressively leaving the SJ domain (Fig. 1B’, between t90 and t115). As the newly assembled SJs acts as a conveyor belt that triggers the basal displacement of the midbody in ECs (Daniel et al., 2018), the difference in midbody basal displacement between EC and SOP predicts a faster SJ remodeling in SOP that we next investigated.

### Remodeling of SJ coincides with SOP midbody basal displacement

We then follow the SJs present in the finger-like protrusions, i.e. the junctions that were present prior to entry into mitosis (Daniel et al., 2018), and discriminate the SJ components of the dividing cell from those of the neighbors. For this purpose, we generated clones of dividing EC or SOP cells expressing MyoIIRFP and no ATPαGFP, but expressing endogenous untagged ATPα, and imaged adjacent to cells expressing ATPαGFP and MyoIIRFP (Fig. 2A, Fig. 2-Fig. S 2A). At t10, the presence of ATPαGFP signal in the finger-like protrusions pointing towards the midbody is originating from the neighbor cell, confirming that the integrity of the SJ barrier is preserved between the SOP and its neighboring EC, as we had previously described between two ECs (Fig. 2A, Fig. 2-Fig. S 2A, SJ plane and (Daniel et al., 2018)). In SOP, as the midbody underwent its basal displacement, at t70±7, the finger-like protrusions thinned and both edges were no longer resolved by light microscopy (t65, Fig. 2A; SJ plane, n=4). At t97±4.5, the finger-like protrusion was barely detectable (t100, Fig. 2A, SJ plane, n=5). Then, the midbody was no longer seen within the plane of the SJs (t110, Fig. 2A, and t115, Fig.1C), prior to be displaced at distance, basal to pIIa nucleus that we interpret as the release of SOP midbody (see below) (t120, Fig. 2A; see also Fig. 1C-C’). In contrast, in EC, the finger-like protrusions connected to the midbody are persisting after t120 (Fig. 1, Fig. S1A and (Daniel et al., 2018)), indicating that the disengagement of finger-like protrusions and midbody release occurs faster in SOP daughters than in ECs.

**Fig. 2.**
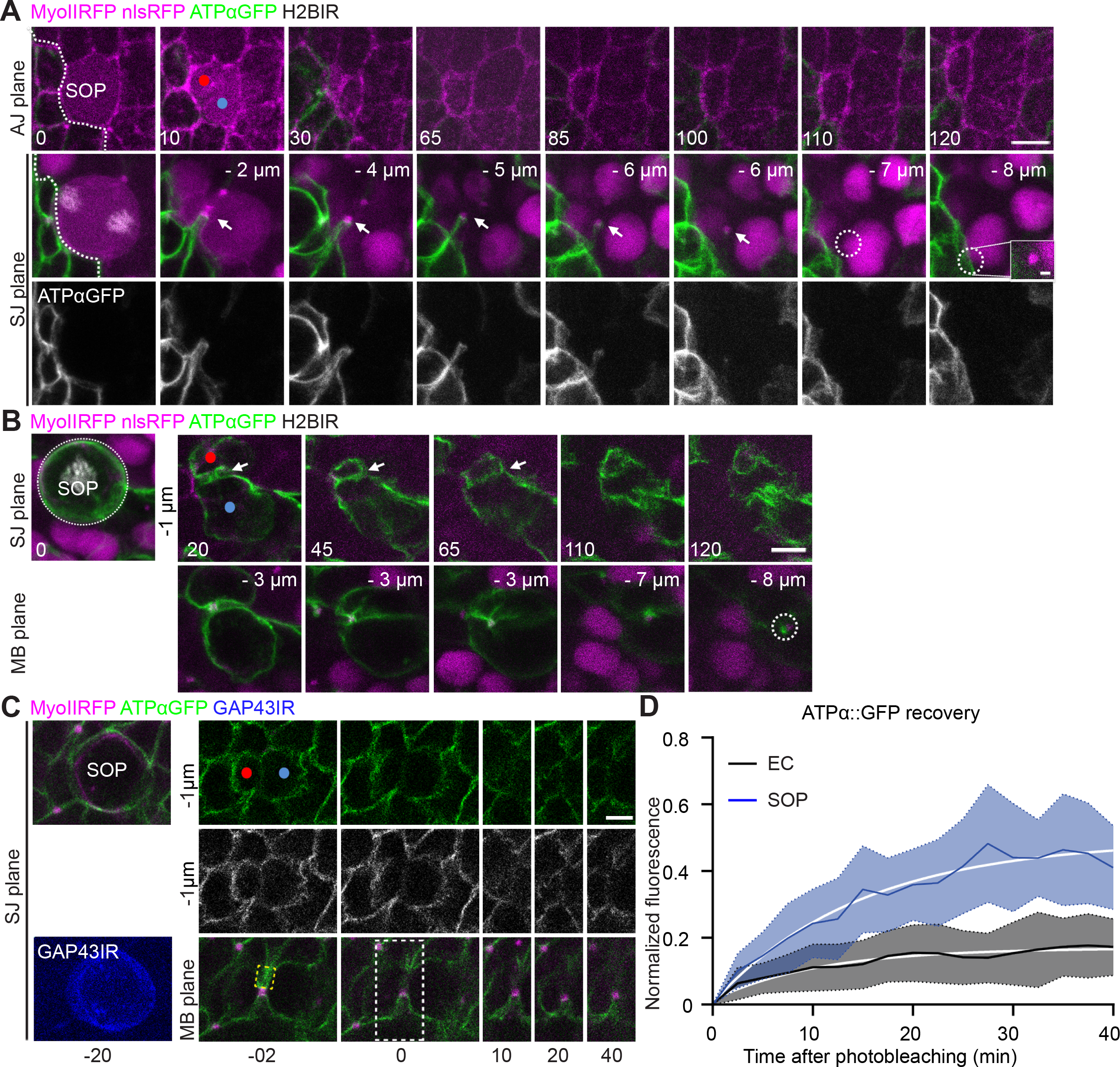
Kinetics of SJ assembly in SOP and EC. (A) Time lapse of SOP (n=5) dividing cells expressing MyoIIRFP and nlsRFP (magenta) and H2BIR (white) adjacent to cells expressing MyoIIRFP (magenta) and ATPαGFP (green and grey on upper and bottom panels, respectively). Distances in μm represent the position relative to the AJ plane determined by apical MyoII signal. White dashed line delineates the clone border at t0. White arrows point to the midbody. Dashed white ring encircles the midbody at t110 and t120. The white square at t120 represents a higher magnification of the midbody located below a nucleus expressing nlsRFP (scale bar represents 1 μm). (B) Time lapse of dividing SOP (n=2) expressing MyoIIRFP (magenta), H2BIR (white) and ATPαGFP (green) close to cells expressing MyoIIRFP (magenta) and nlsRFP (magenta). Dashed white ring encircles the midbody at t120. Distances in μm represent the position relative to the AJ plane determined by apical MyoII signal. (C) Time lapse of SOP dividing cell expressing MyoIIRFP (magenta), ATPαGFP (green and grey on upper and second panels, respectively) and the plasma membrane marker GAP43IR (blue), and FRAP of ATPαGFP. Yellow dotted line square inset indicates the photobleached region of interest. The white dotted line rectangle at t=0 delineate the ROI shown in the right panels (t=10-40 min) at the level of SJ and MB to monitor the recovery of ATPαGFP signal over time. Photobleaching was done 20 min after the anaphase onset with t=0 corresponding to the time of photobleaching. (D) Plot of the quantification of fluorescence recovery of ATPαGFP signal after photobleaching over time in EC (n=20, black) and SOP (n=13, blue) in the SJ plane located 1 μm below AJ. Solid white lines represent a simple exponential fit. AJ: Adherens junctions, SJ: Septate junctions, MB: Midbody. In A, B, and C, SOPs and daughters were identified using H2BIR in A and B (grey) or GAP43IR in C (blue). Red and blue dots mark the pIIb and pIIa respectively (A-C). Time is in minutes with t0 corresponding to the onset of SOP anaphase, except in (C, D) where t0 corresponds to the time of photobleaching. Scale bars represent 5 μm (A-B) and 3 μm (C).

We next analyzed the converse situation to monitor the *de novo* assembly of SJs at the new SOP daughter interface by live-imaging SOP expressing ATPαGFP surrounded by ECs expressing endogenous untagged ATPα. As the cell divides, we observed the appearance of an ATPαGFP signal apical to the midbody (t20, Fig. 2B, SJ plane) which then formed a continuous line, i.e. the novel SJ at t45 (t45, Fig. 2B, SJ plane) compared to t80 in ECs (data not shown and (Daniel et al., 2018)). These results suggest that SJ assembly is faster in SOP daughters than in EC daughters. To confirm this, we performed fluorescence recovery of ATPαGFP signal after photobleaching (FRAP) experiments (Fig. 2C). 40 min after photobleaching one of the two finger-like protrusions of dividing ECs, ATPαGFP signal was barely recovered, presenting mainly an immobile fraction (ymax= 17 % and a t1/2 = 7.4 min, in agreement with previous observations (Fig. 2D, (Daniel et al., 2018)). In contrast, following FRAP of the SOP daughter interface, identified with the plasma membrane marker GAP43∷IRFP670, about 50% of ATPαGFP signal was recovered 40 min at the new daughter interface above the midbody (ymax = 49 % and t1/2 = 10.2 min, Fig. 2C; SJ plane and Fig. 2D).

These data show that the formation of the new SJ, the disengagement of the SJ finger-like protrusions and midbody release occur earlier in SOP daughters than in EC daughters.

### Cytoplasmic isolation and midbody release following SOP division are stereotyped and temporally decoupled

In order to determine when the cytoplasm of the pIIa and pIIb cells are physically separated, we used the photoconvertible probe KAEDE as previously described (Daniel et al., 2018). When photoconverted in the pIIa (or pIIb) cell, photoconverted KAEDE is expected to freely diffuse into the pIIb (or pIIa) cell prior to cytoplasmic isolation, whereas it should be restricted to pIIa (or pIIb) cell after cytoplasmic isolation (Fig. 3-Fig. S3A and data not shown). We used *pnr-*Gal4 to express KAEDE and to subsequently proceed to UAS-GAL4 driven gene silencing.

**Fig. 3.**
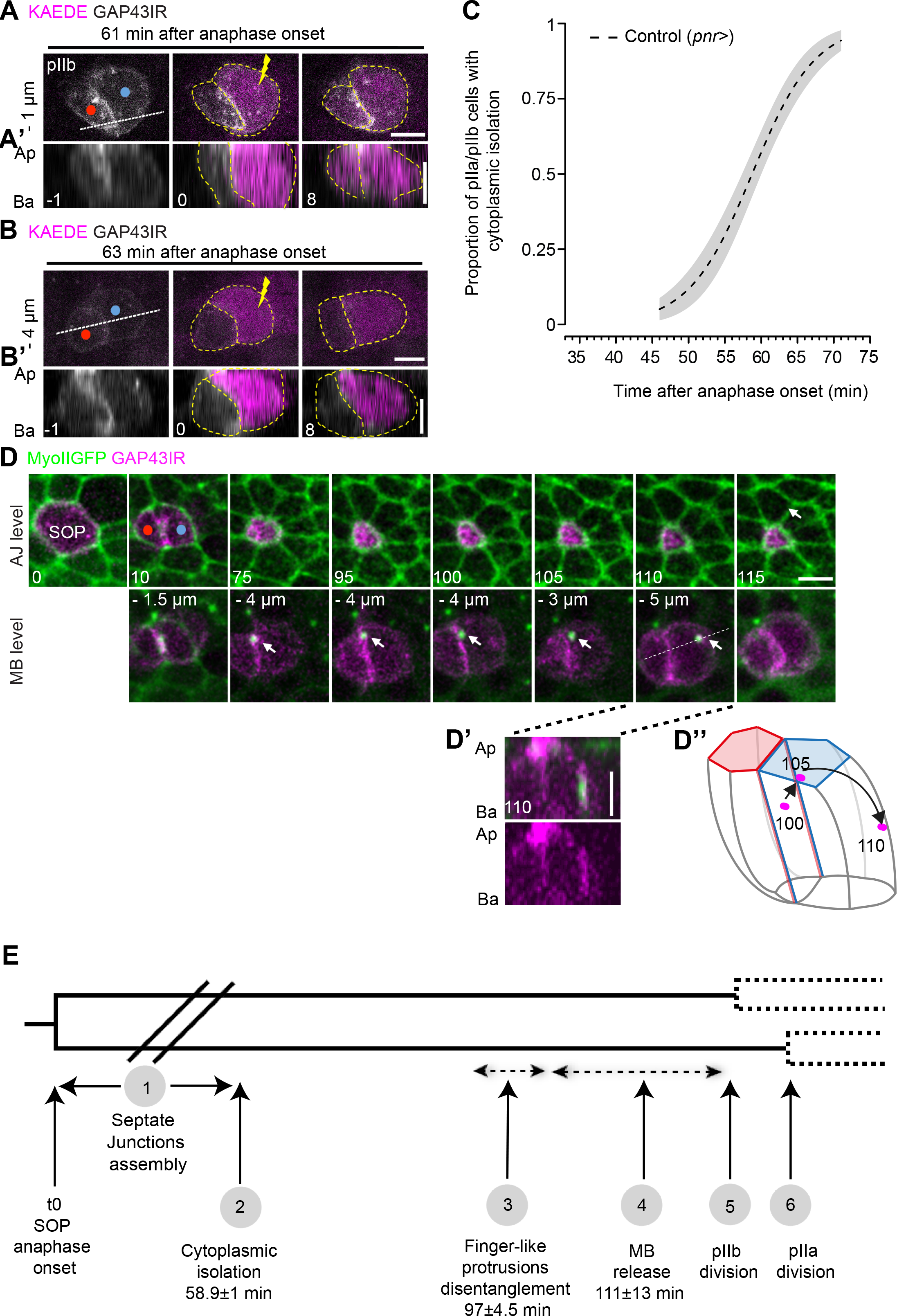
cytoplasmic isolation of pIIa-pIIb and midbody release are temporally controlled. (A-B) Time lapse of SOP dividing cell expressing KAEDE under *pannier* driver. Green to Red (magenta) photoconversion was performed (yellow lightning at t=0) in the pIIb cell at 61 min (A) or 63 min (B) after the onset of anaphase. A’ and B’ correspond to the orthogonal views along the white dashed lines depicted at t=−1 min on panels A and B, respectively. Ap: Apical, Ba: Basal. Time is in min where t=0 corresponds to the time of photoconversion. (C) Plot representing the proportion of pIIa-pIIb cells to have cytoplasmic isolation over time after anaphase onset in control (*pnr*>). Cytoplasmic isolation was assessed based on the ability of photo-converted KAEDE in pIIa/pIIb cell to diffuse in the pIIb/pIIa cell at different time point after the onset of anaphase. The dashed line represents mean values, predicted from a generalized linear model (GLM). SE of the estimates are represented in grey shading. n=62, 24 pupae. (D) Time lapse of SOP dividing cell expressing MyoIIGFP (Green) and GAP43IR (Magenta). White arrows point to the midbody. Distances in μm represent the position relative to the AJ plane determined by apical MyoII signal. (D’) corresponds to the orthogonal view depicted in (D, t110). (D’’) scheme representing the displacement of the midbody over the pIIa membrane at the indicated time as in (D) and adapted from (Houssin et al., 2021). The red and blue cells correspond to the pIIb and pIIb respectively. Ap: Apical; Ba: Basal. (E) Timeline of events during SOP cytokinesis AJ: Adherens junctions, MB: Midbody. Red and blue dots mark the pIIb and pIIa respectively (A, B, D). Time is in minutes with t0 corresponding to the onset of anaphase (D) except in A and B where t0 corresponds to the time of photoconversion. Scale bars represent 5 μm (A-B,D-D’).

For each SOP, identified using GAP43IR, photoconversion of KAEDE was conducted at a specific time point after the onset of anaphase. For example, when photoconversion was performed in the pIIa 61 min after the onset of anaphase, the photo-converted probe diffused into the pIIb cell (Fig. 3A-A‘). By contrast, when photoconversion was performed 63 min post anaphase onset, KAEDE remained restricted in the pIIa cell indicating that cytoplasmic isolation had occurred (Fig. 3B-B‘). This experiment was repeated over several time points ranging from 46 to 71 min (Fig. 3-Fig. S3B), and the result of each individual experiment, i.e. KAEDE able to diffuse or not, were modeled via logistic regression, using the generalized linear model (GLM) (Fig. 3C, see method section). The proportion of cells exhibiting cytoplasmic isolation increased as function of time (Fig. 3C). In our study, due to lethality with the *pnr-*Gal4 driver in one of our conditions, we had to use *scabrous* (*sca-*GAL4 (Mlodzik et al., 1990)). As in the control *pnr*-GAL4 line, we noticed that proportion of cells exhibiting cytoplasmic isolation increased as function of time, but a delay was observed (Fig. 3-Fig. S3B-C and analysis of deviance Table S1A). We estimated the time at which 50% of the pIIa/pIIb cell indicated cytoplasmic isolation (herein termed t1/2, see method section) and determined that t1/2 are 58.9±1.14 min and 66.54 ±1.36 min for *pnr*-Gal4 and *sca*-Gal4 respectively. The reason for the t1/2 difference is unknown between the two conditions and is probably related to the genetic background. For the sake of simplicity in the comparison of effect of gene silencing (see below), since both Gal4 driver lines were used as control values, we arbitrary set the t1/2 of control *pnr*-Gal4 and control *sca*-Gal4 to a relative time centered to 0.

To determine if the cytoplasmic isolation is coupled to the physical cut and release of the midbody, we next monitored the position of the midbody relative to the plasma membrane of the SOP daughter cells, we imaged MyoIIGFP together with the membrane marker GAP43IR (Fig. 3D). 10 min after anaphase onset, the midbody colocalized with GAP43IR at the pIIa-pIIb daughter cells interface (t10, Fig. 3D, MB level) and remained at the pIIa-pIIb interface until t100 (t100, Fig. 3A, MB level). At t105, the midbody leaves the pIIa-pIIb daughter membrane interface (Fig. 3D, MB level, 110.8±12.9 min, n=12), just prior to or concomitant with the pIIb cell anaphase onset (123.4±6.9 min, n=13). After its departure from the pIIa-pIIb interface, the midbody is mainly inherited by the pIIa cell (t105, t110, Fig. 3D, MB level, 91% of the cases, n=11) where it remains attached to the plasma membrane (Fig. 3D’) before the MyoIIGFP signals becomes undetected in pIIa/pIIb cells at t115 (Fig. 3D, 121.5±14 min, n=13). Based on the Fig. 3D, we illustrated the midbody displacement at the pIIa membrane at each time point (Fig. 3D’’). In isolated cells, midbodies can be internalized/phagocytosed in acidic intracellular compartments following abscission (Crowell et al., 2014). To test if the loss of MyoIIGFP signal reflects the loss of GFP signal due to signal quenching by low endosomal pH (Couturier et al., 2014) or the physical disappearance of the midbody, we co-imaged MyoIIRFP and MyoIIGFP together with GAP43IR (Fig. 3-Fig. S3D). Both signals disappeared simultaneously from the pIIa cell and are found outside of the daughter cells at t0 with some GAP43IR signal (Fig. 3-Fig. S3D-D’). We also live-imaged the kinesin-like protein Pavarotti (PavGFP), a member of the centralsplindin complex, together with MyoIIRFP and GAP43IR (Fig. 3-Fig. S3E) and observed that PavGFP and MyoIIRFP signals disappeared simultaneously outside of the daughter cells at t0 with some GAP43IR signal (Fig. 3-Fig. S3E-E’). These data argue that the midbody is not internalized/phagocytosed in the pIIa cell. Instead, the midbody was found at distance from the pIIa cell and was still marked by the GFP, RFP markers as well as the plasma membrane marker GAP43IR indicating that it was in the extracellular space, not in endocytic compartments of epidermal cells (Fig. 3-Fig. S3D’, S3E’).

The sequence of cytoplasmic isolation, the formation of the new bSJs, the disengagement of the SJ finger-like protrusions, and the midbody release in SOP are depicted on a chronological timeline (Fig. 3E). Having described the events and established the timeline, we next investigated the molecular mechanisms underlying epithelial cell abscission of SOP daughters.

### ESCRT III/Shrub localizes at the midbody and in the finger-like protrusions

Based on the evolutionarily conserved function of ESCRT-III/CHMP4B in abscission from archea to vertebrates, we investigated Shrub function in SOP abscission. We first monitored the localization of Shrub. Because available anti-Shrub antibodies give no obvious membrane associated signal following chemical fixation (data not shown; (Pannen et al., 2020)), we made use of a GFP tagged version of Shrub engineered by CRISPR/Cas9 (ShrubGFP, (Mathieu et al., 2022)). We then live-imaged MyoRFP and ShrubGFP (Fig. 4A). t30-60 after the onset of anaphase, a faint signal of ShrubGFP is detected in the plane of the midbody and is slightly enriched along the new daughter cell interface/finger-like protrusion (Fig. 4A’, and 4A’’). At t70±10, while the intensity of the ShrubGFP signal along the pIIa-pIIb interface/finger-like protrusions decreased, ShrubGFP signal was detected in the form of two punctae, one on each side of the midbody, a signal that was still present at t120 (Fig. 4A’, 4A’’’, n=16).

**Fig. 4.**
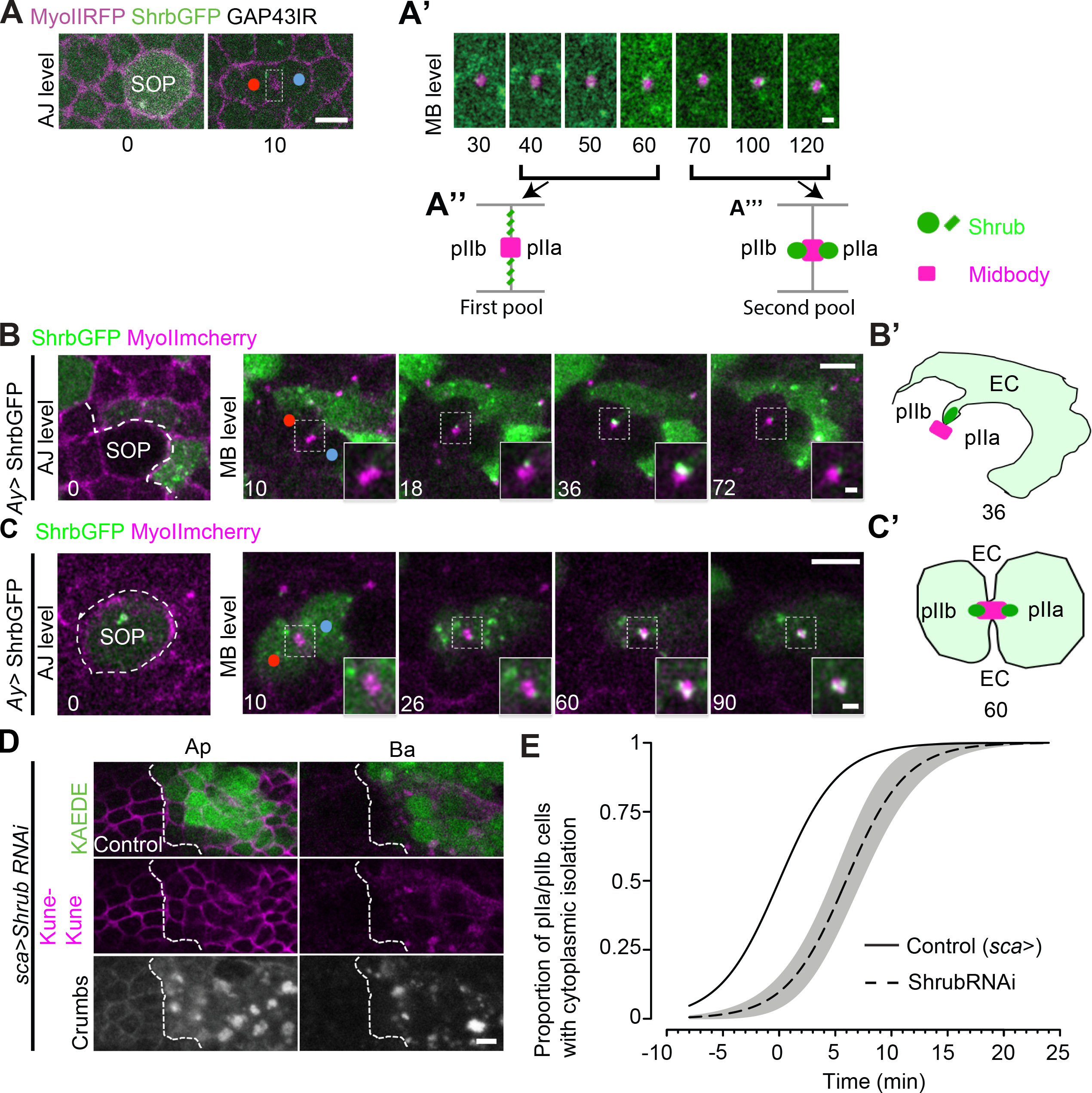
Recruitment and function of Shrub in SOP daughter cells. (A-A’) Time lapse of SOP dividing cells expressing MyoIIRFP (magenta), ShrubGFP (green) and GAP43IR (grey). (A’) Higher magnification of the white dashed line inset depicted in (A) at t10 showing the recruitment of ShrubGFP (green) at the midbody (magenta) over time. (A’’) Schematic representation depicting ShrubGFP recruitment at the presumptive finger like protrusions (between 40 and 60 min) and (A’’’) at the midbody (magenta square) (between 70 and 120 min) corresponding to (A’). (B) Time lapse of SOP dividing cell expressing MyoIImcherry (magenta) adjacent to epidermal cell expressing MyoIImCherry (magenta) together with ShrubGFP expressed under *Ay* Gal4 driver (green). White line delineates the border of clones of cells expressing ShrubGFP at t0. The higher magnifications at the bottom right (MB level, t10-72) correspond to the ROI in the white dotted line square of the corresponding panel (scale bar represents 1 μm). (B’) Representation of the SOP at t36 (B) surrounded by one EC positive for ShrubGFP (light green) and showing the ShrubGFP puncta (dark green) at the tip of the finger-like protrusion pointing toward the midbody (magenta square). (C) Time lapse of dividing SOP expressing MyoIImcherry (magenta) together with ShrubGFP expressed under *Ay* Gal4 driver (Green) close to cells expressing MyoIImcherry (magenta) but not ShrubGFP. White dashed line delineates the clone border at t=0. The higher magnifications at the bottom right (MB level, t10-90) correspond to the ROI in the white dotted line square of the corresponding panel (scale bar represents 1 μm). (C’) Representation of the ShrubGFP positive SOP (light green) at t60 (C) surrounded by ECs negative for ShrubGFP and showing the ShrubGFP punctae (dark green) recruitment at the midbody (magenta square). (D) Localization of KAEDE (green), Kune Kune (magenta) and Crumbs (grey) in nota expressing Shrub RNAi and KAEDE. The white dashed lines separate control from Shrub RNAI (KAEDE expressing cells). Ap: apical, Ba: basal. (E) Plot representing the proportion of pIIa-pIIb cells to have cytoplasmic isolation over time after anaphase onset. Cytoplasmic isolation was assessed based on the ability of photo-converted KAEDE in pIIa/pIIb cell to diffuse in the pIIb/pIIa cell at different time point after the onset of anaphase in control (*sca>*, solid line, n=30, 11 pupae) and ShrubRNAi (*sca>* ShrubRNAi, dashed line, n=30, 14 pupae). Lines represent mean values, predicted from a GLM of t1/2 values for *sca*> centered to 0; SE of the estimates are represented in grey shading. p-value=0,003, **)

To determine if the ShrubGFP signal adjacent to the midbody was coming from the SOP, the neighbors or both (Fig. 4A’), we imaged clones of SOP and EC dividing cells expressing MyoIIRFP devoid of ShrubGFP close to cells expressing ShrubGFP under the *Ay-*Gal4 system (Fig. 4B, Fig. 4-Fig. S4A). A ShrubGFP-positive punctum emanating from the non-dividing neighboring cell was detected at the tip of the finger-like protrusion pointing to the midbody of the SOP (from t18 to t72, Fig. 4B and t36, Fig. 4B’) or to the midbody of the EC (Fig. 4-Fig. S 4A and t36 Fig. 4-Fig. S4A’). In the converse situation in which clones of SOP dividing cells expressing ShrubGFP were close to cells devoid of ShrubGFP (Fig. 4C), at t26 a first punctum of ShrubGFP was appearing at the midbody before the appearance of a second punctum at t60 (Fig. 4C and Fig. 4C’). At t90, the punctae were still present on both side of the midbody of the SOPs (Fig. 4C). In EC, a punctum is detected on one side of the midbody from t28 to t90 (Fig. 4-Fig. S4B and t60 Fig. 4-Fig. S4B’). In contrast to SOPs, we never observed punctae of ShrubGFP on each side of the midbody in the EC. These data revealed the existence of two pools of Shrub, one in the finger-like protrusion and one at the midbody of the SOP, calling for two distinct functions of Shrub.

### Shrub regulates the timing of cytoplasmic isolation of SOP daughter cells

Based on the localization of ShrubGFP at the midbody, we next investigated if Shrub was involved in the abscission by interfering with Shrub function. However, Shrub is required in several processes making difficult to analyze its function. Indeed, clones of epithelial cells homozygous mutant for a null allele of *Shrub*^G5^ delaminate even when apoptosis was suppressed by p35 expression (Hay et al., 1994), preventing us from analyzing abscission in *Shrub* homozygote mutant cells. To circumvent this problem, we opted for an UAS/Gal4-based tissue inducible-based RNAi approach using *sca-*GAL4. We report that silencing of Shrub, in areas where shrub is highly depleted, caused the enlargement of HRS-positive endosomes (Fig. 4-Fig. S4C), the accumulation of Crumbs in endocytic compartments as reported in the *Drosophila* trachea (Dong et al., 2014), and the accumulation of the Claudin-like Kune Kune in intracellular compartments (Fig. 4D) as reported for other claudin-like proteins (Pannen et al., 2020), NrxIV as well as sinuous (Fig. 4-Fig. S4D-E). These three phenotypes are characteristic by the impact of loss of ESCRT-III on intracellular trafficking (Dong et al., 2014; Vaccari et al., 2009), attesting to the effectiveness of the silencing approach. We next determined the consequence of Shrub depletion on the timing of cytoplasmic isolation using the KAEDE photoconversion assay. We performed our experiments in areas in which KAEDE is expressed only in SOP and daughters. In this part of the notum, this mild depletion of shrub does not noticeably affect the distribution of SJ components (data not shown). We found that SOP-specific silencing of Shrub caused a delay in cytoplasmic isolation (Fig. 4E, with a t1/2 delayed of 5.9±1.4 min compared to the *sca* control and Table S1B) indicating that Shrub regulates the timing of cytoplasmic isolation. This data identifies a cell-autonomous function of Shrub in abscission.

We also observed that ShrubGFP is a mutant allele of Shrub, causing a delay in abscission in the female germline stem cyst of *Drosophila* heterozygote for ShrubGFP (67/ 181 GSC with delayed abscission, Fig. 5-Fig. S5A). Similarly in the notum, we observe that the presence of one copy of ShrubGFP delayed the timing of cytoplasmic isolation (Fig. 5, t1/2 +10.3±1.4 min compared to *pnr* control, Table S1C). This delay in abscission observed with shrubGFP is also likely to explain the delay between the detection of two punctae of ShrubGFP at the midbody (Fig. 4A’, t=70±10 min) and the timing of abscission measured in control cells devoid of ShrubGFP (Fig. 3E, t1/2= 58.9±1.1min). Although ShrubGFP is an allele of Shrub, ShrubGFP signal observed on both sides of the midbody is specific as CHMP2BGFP (charged multi vesicular body protein 2b tagged GFP), another ESCRT-III component recruited in a Shrub dependent manner (Babst et al., 2002), exhibits a similar localization pattern (Fig. 5-Fig. S5B). We conclude that Shrub is recruited in a cell autonomous manner on both side of the SOP midbody to control cytoplasmic isolation.

**Fig. 5.**
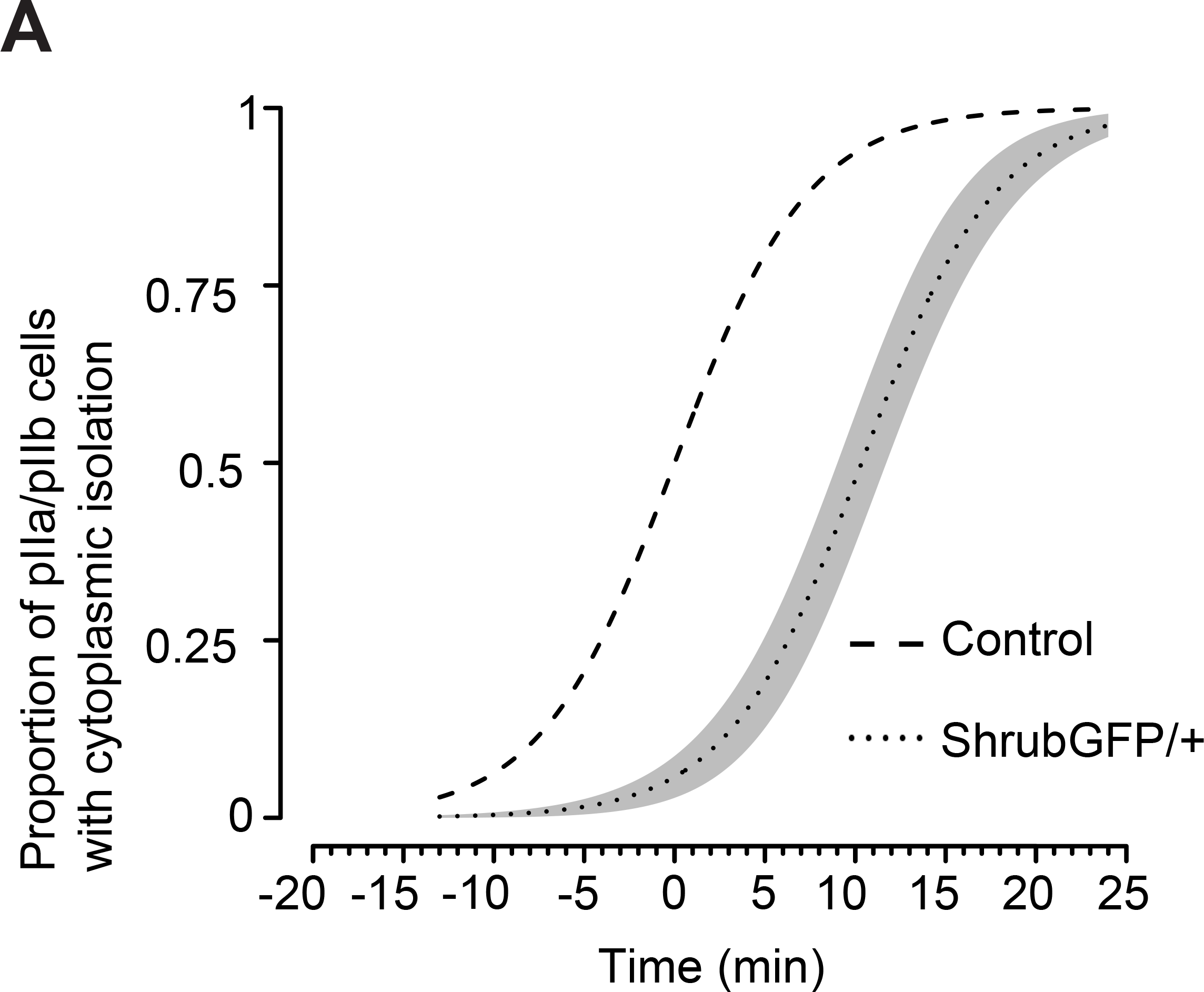
ShrubGFP is a mutant allele that impacts the timing of cytoplasmic isolation. (A) Plot representing the proportion of pIIa-pIIb cells to have cytoplasmic isolation over time after anaphase onset. Cytoplasmic isolation was assessed based on the ability of photo-converted KAEDE in pIIa/pIIb cell to diffuse in the pIIb/pIIa cell at different time point after the onset of anaphase in control (*pnr* >, dashed line, n=62, 24 pupae) and ShrubGFP/+ (*pnr* >+ ShrubGFP/+, dotted line, n=52, 12 pupae). Lines represent mean values, predicted from a GLM of t1/2 values for *pnr*> centered to 0; SE of the estimates are represented in grey shading (p-value=1,68.10^−6^,***). Time is in minutes.

### Coordination between bSJs assembly and timing of abscission

The fact that Shrub is recruited in the finger-like protrusion at a location where SJ remodeling is taking place, and the fact that Shrub depletion has an impact on the integrity of the SJs, prompted us to next investigate whether SJ components could influence the timing of cytoplasmic isolation. We found that depletion of Cora in the SOP and the ECS causes a premature cytoplasmic isolation (Fig. 6A, B; with a t1/2 - 5.8±1.7 min compared to t1/2 *pnr* reference control and Table S1C). We also tracked the position of the midbody upon Cora depletion by live-imaging MyoIIGFP together with GAP43IR (Fig. 6C). 10 min after anaphase onset, the midbody co-localized with GAP43IR at the interface between the two daughter cells (t10, MB level, Fig. 6C). Then, the midbody is released from the interface at t70 (Fig. 6C, MB level, 88.7±14.8 min, n=8) and is not detected within the pIIa cell at t85 (Fig. 6C, 94.3±11.3 min, n=7). Thus, the midbody release and its disappearance from the pIIa cell happen earlier than in the control (Fig. 6D and 6E respectively). This is not caused by an acceleration of cell cycle in Cora depleted cells as the entry into anaphase onset of pIIb and pIIa takes place later compared to the control (Fig. 6F). Together, these data shows that abscission happens earlier upon Cora depletion than in the control and indicates that the formation of SJs delays abscission timing of the SOP.

**Fig. 6.**
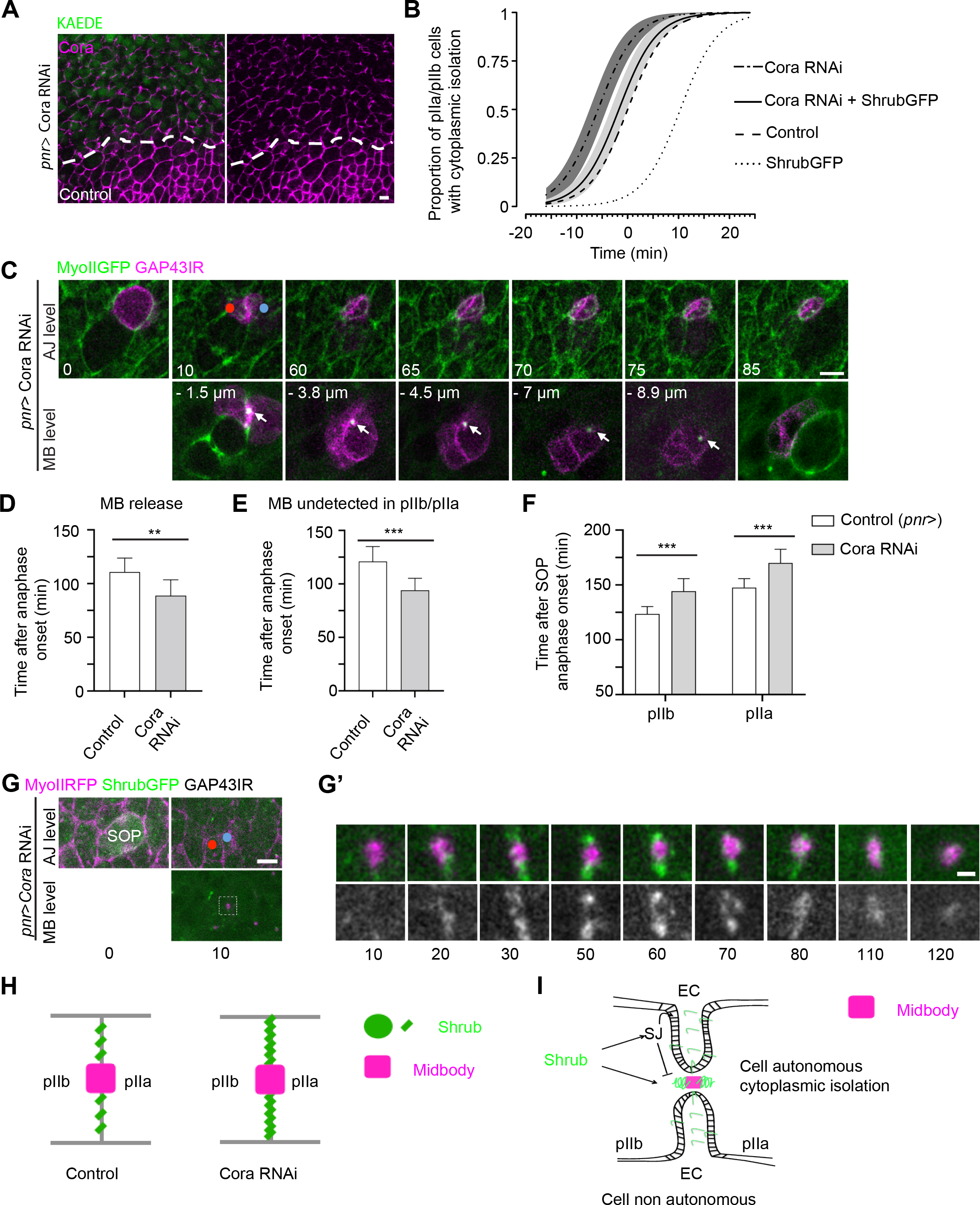
Interplay between Shrub and SJ components in the control of cytoplasmic isolation. (A) Localization of Cora (magenta) upon silencing of Cora (A). The white dashed lines separate wild type from KAEDE expressing cells, which correspond to the area of Cora depletion. (B) Plot representing the proportion of pIIa-pIIb cells to have cytoplasmic isolation over time after anaphase onset. Cytoplasmic isolation was assessed based on the ability of photo-converted KAEDE in pIIa/pIIb cell to diffuse in the pIIb/pIIa cell at different time point after the onset of anaphase in control (*pnr>*, dashed line, n=62, 24 pupae), Cora RNAi (*pnr*>CoraRNAi, dotdashed line, n=32, 16 pupae), ShrubGFP/+ (*pnr*>+ ShrubGFP/+, dotted line, n=52, 12 pupae) and Cora RNAi + ShrubGFP/+ (*pnr*>CoraRNAi+ShrubGFP/+ solid line, n=30, 6 pupae). Lines represent mean values, predicted from a GLM of t1/2 values for *pnr*> centered to 0; SE of the estimates are represented in grey shading. (*pnr> vs pnr*>CoraRNAi, p-value=0,029,*; *pnr> vs* Cora RNAi + ShrubGFP/+, p-value=0.425, ns; Cora RNAi + ShrubGFP/+ vs *pnr*>+ ShrubGFP/+, p-value=4,52.10^−7^,***; *pnr*>CoraRNAi vs Cora RNAi + ShrubGFP/+, p-value=0.075,ns). (C) Time lapse of SOP dividing cell depleted of Cora expressing MyoIIGFP (Green) and GAP43IR (Magenta). White arrows point to the midbody. Distances in μm represent the position relative to the AJ plane determined by apical MyoII signal. (D) Plot representing the time after anaphase onset (min) when the midbody is released from the new interface between pIIb and pIIa in control (white, n=12) and Cora depleted cells (light grey, n=8). Time is in min. (unpaired t-tests; p-value=0,002, **) (E) Plot representing the time after anaphase onset (min) when the midbody is disappearing from the SOP in control (white, n=13) and Cora depleted cells (light grey, n=7). Time is in min. (unpaired t-tests; p-value=0.0004, ***) (F) Plot of the time of the onset of pIIa and pIIb anaphase after the onset of SOP anaphase in control (black, n=13) and Cora depleted (grey, n=8) cells. Time is in min. (unpaired t-tests; p-value=0.000045 and p-value=0,00081,*** for pIIb and pIIa respectively) (G) Time lapse of SOPs expressing MyoIIRFP (magenta) and ShrubGFP (green) depleted from Cora. (G’) Higher magnifications of the white dotted square inset depicted in (C) at t10 showing ShrubGFP (green) recruitment to the midbody (magenta) (n= 2/6). (H) Scheme depicting Shrub recruitment (Green) in the finger-like protrusions and at the level of the midbody (magenta rectangle) during cytokinesis in SOP control and depleted with Cora RNAi. (I) Scheme proposing a model representing Shrub functions during epithelial cytokinesis. Shrub exerts a cell autonomous function in the control of the separation of the pIIa-pIIb cytoplasm. Shrub (green spiral) is recruited at the level of the midbody (magenta rectangle). Shrub exerts a cell non-autonomous function in the remodeling of the SJs within the finger-like protrusions. This activity is linked to SJ trafficking via endosomes. Time is in minutes with t0 corresponding to the onset of anaphase (C-E, G-G’). Red and blue dots mark the pIIb and pIIa respectively (C, G). Scale bars represent 5 μm (A, C, G) and 1 μm (G’).

Thus, Shrub and the SJ component Cora act as positive and negative regulators of cytoplasmic isolation of SOP daughters, respectively. However, because depletion of Shrub leads to altered localization of SJ components ((Pannen et al., 2020); Fig. 4D and - Fig. S4D-E) the above results are counterintuitive at a first glance. Indeed, in a simplistic model in which Shrub would only be required for SJ remodeling, depletion of Shrub by causing a reduction in the amounts of SJ components should lead to early cytoplasmic isolation. Nevertheless, as indicated above, Shrub also localizes at the midbody level where it promotes abscission. These data on the dual function of Shrub led us to further study the relationship between Shrub and Cora. Silencing of Cora caused an increased recruitment of ShrubGFP within the finger-like protrusion in one third of the cases, (n=6; Fig. 6G’, t30-t70 and Fig. 6H) further suggesting a function for Shrub at the site of local SJ remodeling (see discussion). Because, ShrubGFP behaves like a mutant allele delaying abscission (Fig. 5E), we reasoned that the presence of ShrubGFP could modify the timing of abscission measured upon simple depletion of Cora. The timing of cytoplasmic isolation upon Cora depletion in the presence of ShrubGFP mutant allele caused a premature isolation compared to ShrubGFP allele alone (t1/2 difference of 12.1 min). Plotting results also suggest a tendency for Cora depleted cells to exhibit greater proportions of a premature cytoplasmic isolation than when shrubGFP allele is expressed in Cora depleted cells (t 1/2 difference of 4.09 min).

Overall, these data indicate that Shrub exerts two functions during SOP cytokinesis: a first function, at the level of the SOP midbody to control the timing of cytoplasmic isolation; and a second cell non-autonomous function in the finger-like protrusions for the SJ-based remodeling (Fig. 6I).

## Discussion

In this study, we further characterized the coordination of abscission and the maintenance of epithelial permeability barrier functions during cytokinesis in dividing SOPs and compared them to symmetrically dividing epidermal cells. In both types of cells, the midbody assembles below the newly forming pIIa-pIIb adherens junctions. In the plane of the midbody, the neighboring cells remains in contact with the divided cell through finger-like protrusions. These finger-like protrusions are located within the layer of SJs that were already connecting the neighbors and the SOP prior to SOP mitosis. The disassembly of SJs appears to be the rate limiting step for the disentangling of the finger-like protrusions. We reported that the basal displacement of the midbody and the *de novo* assembly of SJs occurs faster in SOP than in EC, resulting in a quicker disentanglement of the finger-like protrusions, an event that precedes midbody release. We have described that cytoplasmic isolation is temporally uncoupled to midbody release, suggesting that abscission is a two-step process. At the molecular level, SJ components negatively regulate the timing of cytoplasmic isolation implying a regulatory role of permeability barrier maintenance on abscission. Finally, we reported that Shrub exerts two complementary functions in SOP abscission. A first, cell autonomous control of the timing of cytoplasmic isolation at the level of the SOP midbody. The second function of Shrub is at the finger-like protrusions to remodel SJs and prevent premature abscission until the permeability barrier is fully assembled.

### Dynamics of bSJs assembly and midbody release in SOP compared to EC

FRAP analyses revealed that the percentage of mobile fraction of ATPαGFP is increased in SOPs compared to ECs, suggesting that the assembly of SJs at the new pIIa-pIIb interface occurs faster in SOPs. In addition, the disentanglement of the finger-like protrusions, corresponding to the completion of disassembly of SJs between SOP and neighbors also occurs faster in SOPs. This is associated with a faster basal displacement of the midbody. These results add arguments in favor of the concept that *de novo* assembly of SJs acts as a conveyor belt of midbody basal displacement (Daniel et al., 2018). As the endosomal system actively contributes to transport and turnover of SJ complexes (Nilton et al., 2010; Pannen et al., 2020; Tempesta et al., 2017; Tiklová et al., 2010), our data suggest some differences in the intracellular trafficking between SOP and EC. As SOP daughters enter into mitosis about 90-120 min after the anaphase onset of SOP division, the dynamics of membrane trafficking might be under cell cycle regulation. It is also possible that the transcriptional program of SOPs includes specific membrane trafficking regulators or SJ components that directly impact the kinetics of assembly of SJs. In addition, as the disentanglement of the finger-like protrusions occurs faster in SOPs, our data also implies that the dynamic of assembly and disassembly of SJs in the SOPs and its daughters dictates the dynamics of SJ components in the neighboring ECs. Thus, the dynamics of SJ between ECs and SOPs is different than between two ECs.

### Role of SJ components in regulating SOP cytoplasmic isolation

We report that loss of SJ components results in premature cytoplasmic isolation, raising the question of how can SJ components negatively regulate abscission? In the finger-like protrusions formed as the result of cytokinetic ring constriction, the nascent midbody is embedded in the SJs that were assembled between the SOP and its EC neighbors prior to mitosis. This topology ensures that at the level of the midbody within the finger-like protrusion, the permeability barrier function is maintained. Above the midbody, the novel SJ between SOP daughters then progressively assembles to build the permeability barrier between the daughters. We envision the topology of the finger-like protrusions as a mean of maintaining the mother-daughter permeability barrier throughout cytokinesis. The disentanglement of finger-like protrusions corresponds to the dismantlement of the old SJ leading to the release of the midbody. Thus, apical to basal displacement of the finger-like protrusions involves intense membrane remodeling and probably membrane trafficking which include Shrub-dependent trafficking. Upon Depletion of Cora, Shrub is found at higher levels in the finger-like protrusions, and the process leading to abscission and midbody release is accelerated. While the molecular trigger is yet to be characterized, a simple possibility is that SJ components negatively regulate Shrub recruitment at the level of the midbody and in the finger-like protrusions.

### Temporal uncoupling between cytoplasmic isolation and midbody release

In isolated vertebrate cells, the cytoplasmic isolation is concomitant with abscission, i.e. the physical separation of the divided cells (Guizetti et al., 2011; Steigemann et al., 2009). We showed here, as demonstrated in the C. elegans one cell embryo (Green et al., 2013), that cytoplasmic isolation and midbody release are temporally separated events (Fig. 3F). A possible caveat in our study resides in the use of the KAEDE probe, a 116kDa homotetramer (Ando et al., 2002). Indeed, we cannot exclude that a probe of smaller size or a solute would still be able to equilibrate after the time we have here determined as the timing of cytoplasmic isolation. However, the fact that Shrub is recruited on both side of the midbody and regulates the time when the KAEDE probe is no longer able to diffuse between the divided cells argues that cytoplasmic isolation occurs several minutes prior to midbody release. As the midbody release coincide with the disentanglement of the finger-like protrusions (Fig. 3F), we propose that the midbody is embedded within the SJ strands and released only upon SJ disassembly. Following its release from the pIIa-pIIb interface, the midbody is displaced on the surface of the pIIa cell until it detaches and found at distance from the pIIa cell. There, the midbody is still labelled with MyoIIGFP/PavGFP, MyoIIRFP and GAP43IR markers suggesting that the midbody is in the extracellular space rather than being internalized by adjacent epidermal cells.

### Cell autonomous and cell non-autonomous functions of Shrub

We report that Shrub positively and cell autonomously regulates abscission in SOP as also observed in mammalian cultured cells and *Drosophila* germline stem cell (Eikenes et al., 2015; Matias et al., 2015). ShrubGFP appears in the form of punctiform signal in a stereotyped manner on both sides of the SOP midbody. These two puncta of ShrubGFP are recruited at the midbody long before the midbody release. This is also observed in female *Drosophila* germline stem cells where Shrub is recruited to the ring canal from G1/S prior to abscission in G2 phase of the following cell cycle (Eikenes et al., 2015; Matias et al., 2015). Interestingly, recruitment of Shrub occurs around the S to G2 transition of the daughter cells and about one hour after the entry of SOP into mitosis (Audibert, 2005). Then, the midbody release takes place just before the G2/M transition of the pIIb cell. Thus, the temporal regulation of abscission in SOP appears to follow a similar cell-cycle dependent regulation as the *Drosophila* germline stem cells. Whether this cell-cycle dependence is specific to stem cells and progenitors or also applies to epidermal cells await further investigation.

Another pool of Shrub is acting in a cell non-autonomous manner. As in SOPs, ShrubGFP is also recruited at the midbody of the dividing epidermal cell and is recruited in the finger-like protrusions. What could be the function of the pool of ShrubGFP in the finger-like protrusions? Based on the fact that Shrub regulates SJs steady state distribution and dynamics (Pannen et al., 2020), and is also known to regulate endosomal sorting, Shrub could locally participate in SJ remodeling. Alternatively, based on the geometry of the tip of the finger-like protrusions and possible local constraints linked to basal displacement of the SOP midbody, Shrub could be recruited there to promote membrane repair (Jimenez et al., 2014; Scheffer et al., 2014) and/or locally compensate for leakage in permeability barrier function. Consistent with this model, when the permeability barrier function is challenged, as for example during Cora depletion, ShrubGFP is prone to accumulate in the finger-like protrusions.

### Abscission timing, midbody inheritance and cell fate decision

A link between the regulation of cell division and fate decision is reported and abscission is thought to regulate fate acquisition (Chaigne et al., 2020; Ettinger et al., 2011). Here, we found that the abscission is asymmetric with the midbody remnant inherited by the pIIa cell in most cases. By analogy with the C. elegans single-cell embryo, the midbody could be inherited by default by the cell that will divide later than its daughter (Green et al., 2013). Can SOP midbody remnant signal and impact on proliferation, differentiation and/or cell fate as in mammalian cells (Ettinger et al., 2011; Kuo et al., 2011; Peterman et al., 2019; Pohl and Jentsch, 2009) ? Whether SOP midbody exerts a function on proliferation awaits further monitoring of its behavior (uptake?) after its release from the pIIa cell.

The fact that abscission occurs rather late after the anaphase onset while Notch-dependent fate acquisition is initiated about 15 min after the onset of anaphase raises an intriguing question (Bellec et al., 2018; Couturier et al., 2012). Indeed, our photo-activation experiment shows that pIIa and pIIb share their cytoplasm until t60 after anaphase, which contrasts with the fact that the Notch intracellular domain (NICD) is translocated exclusively in the nucleus of pIIa. The cell fate determinants Numb and Neuralized, two cytosolic proteins, are also unequally inherited by the pIIb cell before cytoplasmic isolation. An interesting question is how NICD, and Numb, Neuralized remain confined to pIIa and pIIb cells respectively, while at the same time the KAEDE probe freely diffuses bidirectionally. This situation is reminiscent to that of phospho-Mad that remains restricted to the most anterior of *Drosophila* germline stem cells in stem cysts (Eikenes et al., 2015; Mathieu et al., 2013; Matias et al., 2015). Unless, Notch signaling components are assembled in very large protein complexes or with cytoskeletal components that prevent them from diffusing through the opened intercellular bridge, these data imply that selective passage must be regulated at the intercellular bridge (Mullins and Biesele, 1977; Norden et al., 2006; Steigemann et al., 2009), as reported for proteins and organelles in yeast (Lengefeld and Barral, 2018).

Overall, our study sheds light on a complex coordination of abscission, maintenance of epithelial permeability barrier functions and fate acquisition in SOPs. Based on the apico-basal topology of mechanical and permeability barriers, apical positioning of the midbody within the tight junctions in vertebrates (Dubreuil et al., 2007; Higashi et al., 2016; Jinguji and Ishikawa, 1992) and the role of ESCRT components in tight junction protein trafficking (Dukes et al., 2011; Raiborg and Stenmark, 2009), it is tempting to speculate that the cell autonomous and non-autonomous effects we described in this study may also be at play during epithelial abscission of progenitors in vertebrates.

## Acknowledgements

We thank Vanessa Auld, Anne Uv, the Bloomington Stock Center, the Vienna *Drosophila* RNAi Center and the National Institute of Genetics Fly Stock Center for providing fly stocks. We also thank S. Dutertre and X. Pinson from the Microscopy Rennes Imaging Center-BIOSIT (France). The monoclonal antibodies against Elav, Cut, Cora, DE-Cad and α-spectrin were obtained from the Developmental Studies Hybridoma Bank, generated under the auspices of the National Institute of Child Health and Human Development, and maintained by the University of Iowa Department of Biological Sciences. We thank Arnaud Echard, Thomas Esmangart de Bournonville, and members of RLB’s lab for critical reading of the manuscript. We also thank R.J. Scott Mc Cairns for the help with the GLM models.

## Author contributions

### Funding

This work was supported in part by the La Ligue contre le Cancer-Equipe Labellisée (R.L.B.) and the Association Nationale de la Recherche et de la Technologie programme PRC Vie, santé et bien-être CytoSIGN (ANR-16-CE13-004-01 to R.L.B.).

## Materials and Methods

### Key resources table

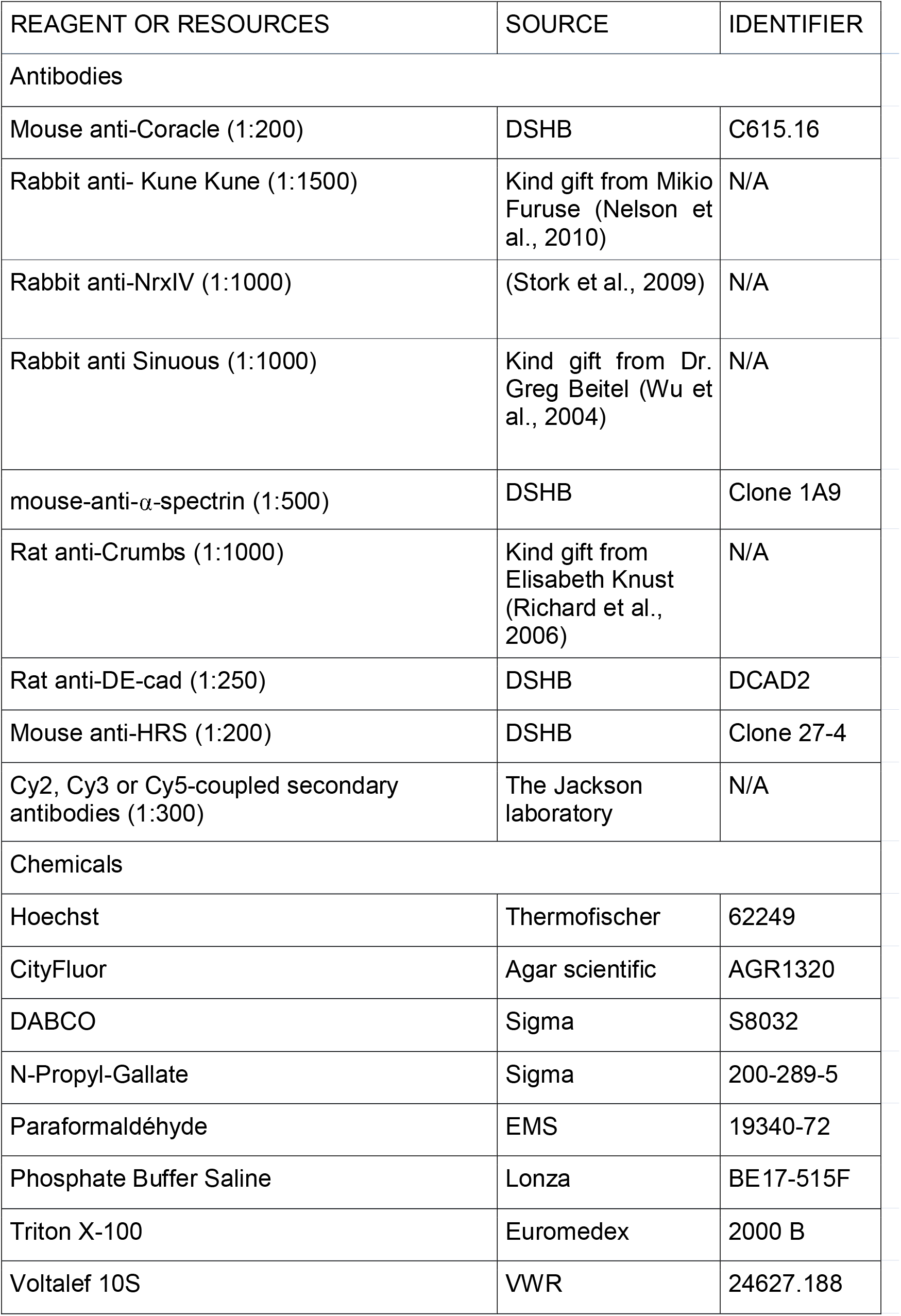
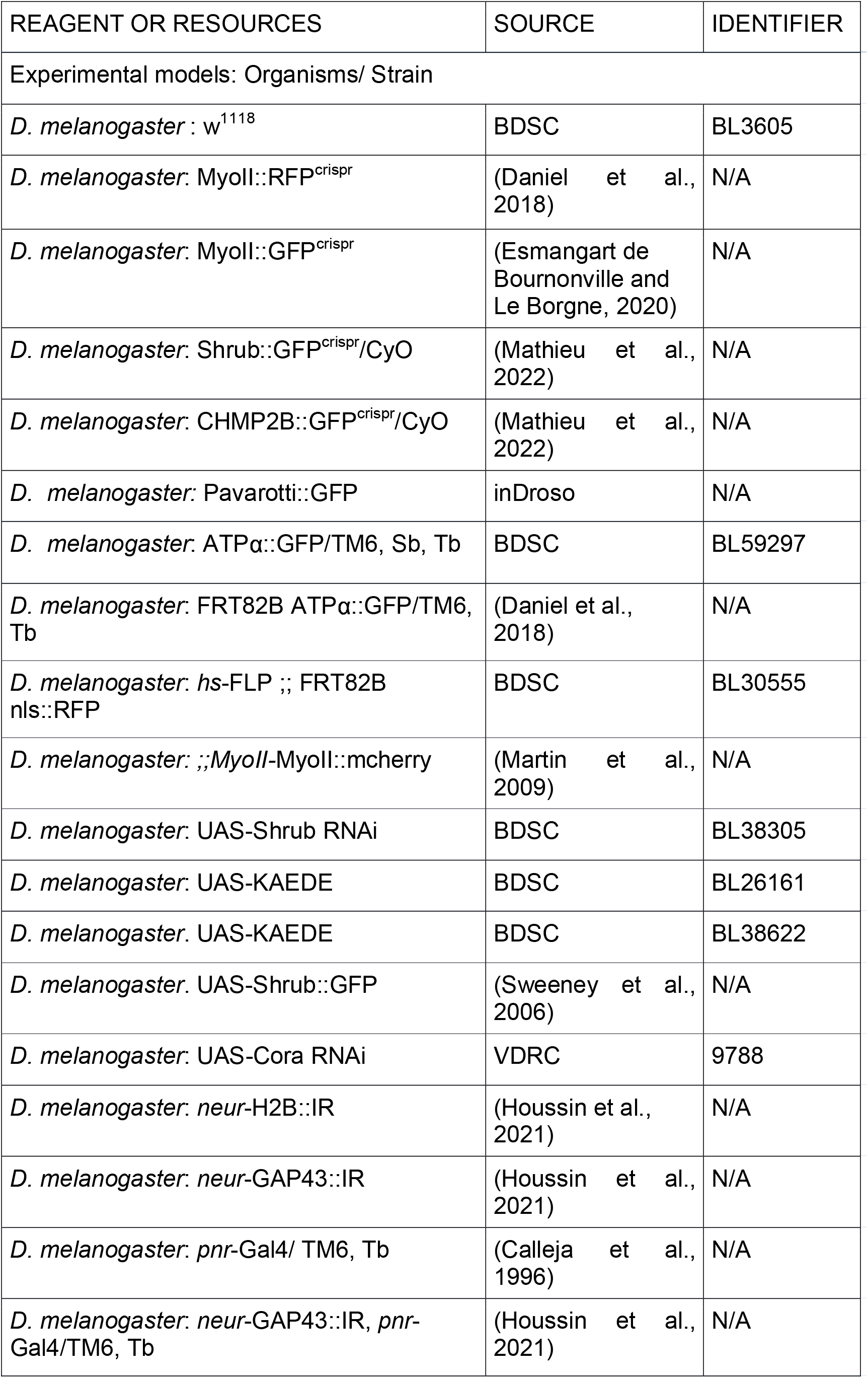
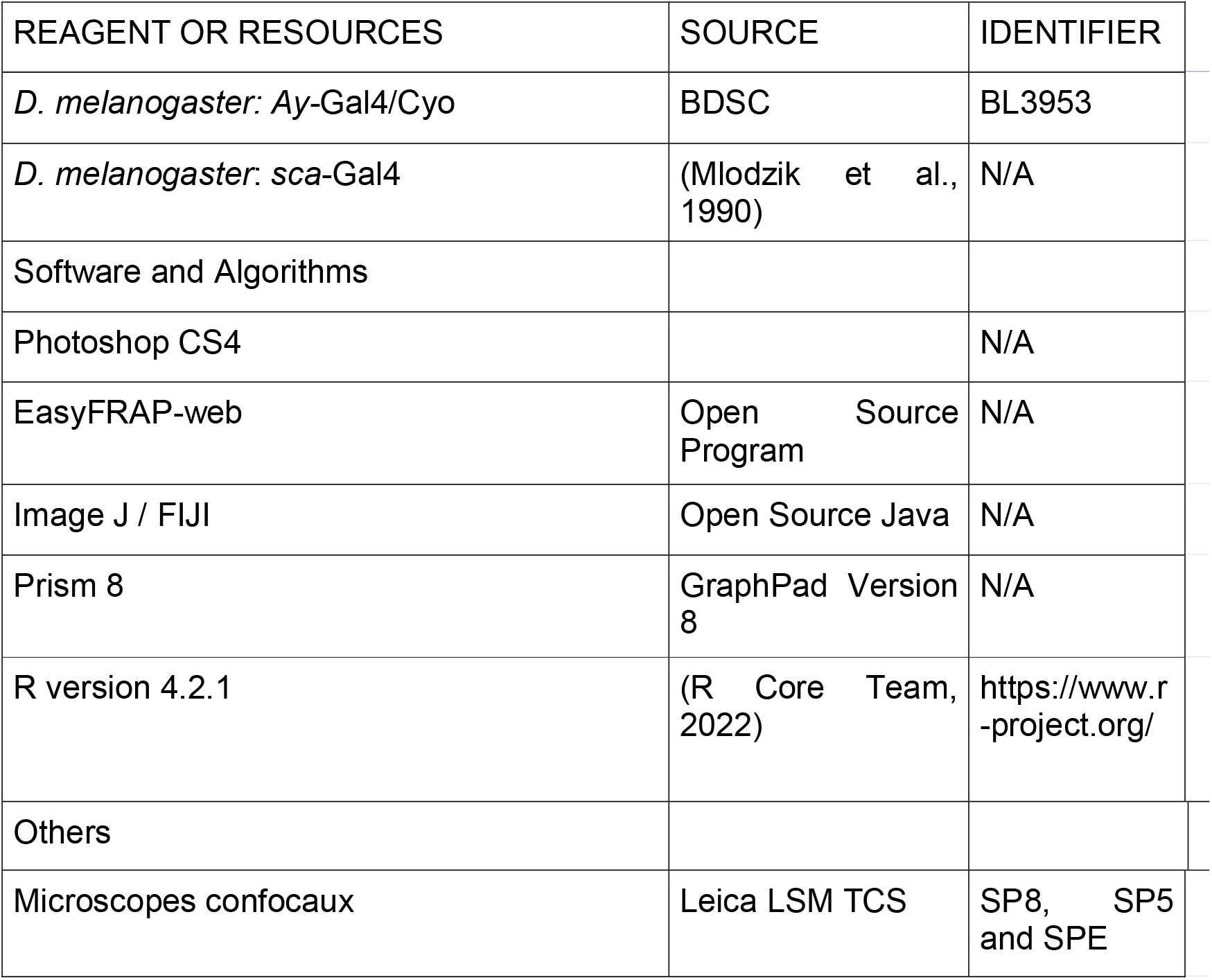

#### Drosophila stocks and genetics

*Drosophila* melanogaster stocks were maintained and crossed at 25°C.

Somatic clones were induced using the *hs*-FLP with two heat shocks (60 min at 37°C) at second and third instar larvae. *pnr*-Gal4 was used to drive the expression of UAS-Cora RNAi and the photoconvertible probe UAS-KAEDE. *Ay*-Gal4 was used to drive the expression of UAS-ShrubGFP. *sca*-Gal4 was used to drive the expression of UAS-ShrubRNAi and UAS-KAEDE.

Declaration of contained use of genetically modified organisms (GMOs) of containment class n°2898 from the French Ministère de l’Enseignement Supérieur, de la Recherche et de l’Innovation.

#### Drosophila genotypes

**Fig. 1**

(B) MyoII∷RFP ; *neur*-H2B∷IR/+ ; ATPα∷GFP/+

(C-C’) *neur*-H2B∷IR

(E) MyoII∷RFP ; *neur*-H2B∷IR/+ (EC and SOP) and MyoII∷RFP ; *neur*-H2B∷IR/+ ; ATPα∷GFP/+ (EC and SOP)

**Fig. 2**

(A-B) *hs*-FLP/MyoII∷RFP ; *neur*-H2B∷IR/+ ; FRT82B ATPα∷GFP/FRT82B nls∷RFP (C-D) MyoII∷RFP ;; ATPα∷GFP/*neur*-GAP43∷IR

**Fig. 3**

(A-C) UAS∷KAEDE; *neur*-GAP43∷IR, *pnr*-Gal4/+

(D-D’) MyoII∷GFP, *neur*-H2B;; *neur*-GAP43∷IR, *pnr*-Gal4/+

**Fig. 4**

(A-A’) MyoII∷RFP ; Shrub∷GFP/+; *neur*-GAP43, *pnr*-Gal4∷IR/+

(B-C) *hs*-FLP; UAS-Shrub∷GFP/*Ay*-Gal4; *MyoII*-MyoII∷mcherry

(D) *sca*-Gal4/UAS-Shrub-RNAi; UAS-KAEDE/+

(E) *sca*-Gal4/UAS-Shrub-RNAi; UAS-KAEDE/+, *sca*-Gal4/UAS-Shrub-RNAi; UAS-KAEDE/*neur*-GAP43∷IR and *sca*-Gal4/+; UAS-KAEDE/+

**Fig. 5**

UAS∷KAEDE; *neur*-GAP43∷IR, *pnr*-Gal4/+ and UAS∷KAEDE/ShrubGFP; *neur*-GAP43∷IR, *pnr*-Gal4/+

**Fig. 6**

(A) UAS-KAEDE/+; *neur*-GAP43∷IR, *pnr*-Gal4/UAS-Cora RNAi

(B) UAS-KAEDE/+; *neur*-GAP43∷IR, *pnr*-Gal4/+ and UAS-KAEDE/+; *neur*-GAP43∷IR, *pnr*-Gal4/UAS-Cora RNAi and UAS-KAEDE/Shrub∷GFP; *neur*-GAP43∷IR, *pnr*-Gal4/+ and UAS-KAEDE/Shrub∷GFP; *neur*-GAP43∷IR, *pnr*-Gal4/UAS-CoraRNAi

(C) MyoII∷GFP, *neur*-H2B;; *neur*-GAP43∷IR, *pnr*-Gal4/ UAS∷Cora RNAi

(D-F) MyoII∷GFP, *neur*-H2B;; *neur*-GAP43∷IR, *pnr*-Gal4/+ and MyoII∷GFP, *neur*-H2B;; *neur*-GAP43∷IR, *pnr*-Gal4/ UAS∷Cora RNAi

(G-G’) MyoII∷RFP; shrubGFP/+; *neur*-GAP43∷IR, *pnr*-Gal4/ UAS∷Cora RNAi

**Fig. S 1**

(A) MyoII∷RFP ; *neur*-H2B∷IR/+ ; ATPα∷GFP/+

**Fig. S 2**

(A) *hs*-FLP/MyoII∷RFP ; *neur*-H2B∷IR/+ ; FRT82B ATPα∷GFP/FRT82B nls∷RFP

**Fig. S 3**

(B) UAS∷KAEDE; *neur*-GAP43∷IR, *pnr*-Gal4/+ and *sca*-Gal4/+; UAS-KAEDE/+

(C) UAS∷KAEDE; *neur*-GAP43∷IR, *pnr*-Gal4/+ and *sca*-Gal4/+; UAS-KAEDE/+

(E) MyoII∷RFP/MyoII∷GFP;; GAP43∷IR/+

(F) MyoII∷RFP;; GAP43∷IR/ Pav∷GFP

**Fig. S 4**

(A,B) *hs*-FLP; UAS-Shrub∷GFP/*Ay*-Gal4; *MyoII*-MyoII∷mcherry

(C-E) *sca*-Gal4/UAS-Shrub-RNAi; UAS-KAEDE/+

**Fig. S 5**

(A) *w^1118^* and Shrb∷GFP/+

(B) MyoII∷RFP ; CHMP2BGFP/+ ; *neur*-GAP43/+

## Methods

### Immunofluorescence

Pupae aged from 16 to 24h after puparium formation (APF) were dissected using Cannas microscissors in 1X Phosphate-Buffered Saline (1X PBS, pH 7.4) and fixed 15 min in 4% paraformaldehyde solution at room temperature (RT) (Gho et al., 1996). Following fixation, dissected nota were permeabilized using 0.1% Triton X-100 in 1X PBS (PBT) and incubated with primary antibodies diluted in PBT for 2 hours at RT. After 3 washes of 5 minutes in PBT, nota were incubated with secondary antibodies diluted in PBT for 1 hour at RT. Following incubation, nota were washed 3 times in PBT, and once in PBS prior mounting in 0,5% N-propylgallate with DABCO dissolved in 1X PBS/90% glycerol.

For *Drosophila* germline stem cell identification, antibody staining and Hoechst staining were performed according to standard protocols. Briefly, ovaries or testis were dissected in PBS, fixed in 4% PFA, rinced and permeabilized in PBT (PBS-0,2%Triton) for 30 min, left overnight with primary antibodies in PBT at 4°C, washed 2 h in PBT, left with secondary antibodies in PBT for 2 hrs at room temperature, washed 1 h in PBT and mounted in Citifluor (Eikenes et al., 2015; Matias et al., 2015).

GSC were identified with the fusome staining (round or linking the CB) and counted. Stem-cysts (more than 2 cells anchored to the niche and linked by a fusome) and polyploid GSC (higher DNA and larger fusome than control) were counted. Stacks were acquired every 0.7 μm.

### Live imaging and image analysis

Live imaging was performed on pupae aged for 16 to 22h APF at 20-25°C (Gho et al., 1999). Pupae were sticked on a glass slide with a double-sided tape, and the brown pupal case was removed over the head and dorsal thorax using microdissection forceps. Pillars made of 4 to 5 glass coverslips were positioned at the anterior and posterior side of the pupae, respectively. A glass coverslip covered with a thin film of Voltalef 10S oil is then placed on top of the pillars such that a meniscus is formed between the dorsal thorax of the pupae and the glass coverslip. Images were acquired with a confocal microscope Leica SP5, SP8 or SPE equipped with a 63X N.A. 1.4. and controlled by LAS AF software. Confocal sections were taken every 0.5 μm unless otherwise specified. All images were processed and assembled using ImageJ/FIJI software and Photoshop CS4.

### Fluorescence recovery after photobleaching

FRAP experiments were performed in pupae expressing MyoIIRFP with ATPαGFP and neur-GAP43IR. Regions of interest corresponding to one of the two ATPα∷GFP finger-like protrusions pointing to the midbody were bleached (488 nm at 100% laser, 1 iteration of 100 ms) using a LSM Leica SP8 equipped with a 63X N.A. 1.4 PlanApo objective. Confocal stacks were acquired every 2 min and 30s after photobleaching.

### KAEDE photoconversion

Photoconversion assays were performed in pupae expressing the green to red photoconvertible probe KAEDE. KAEDE was photoconverted (405 nm laser at 0,5 to 2% power, point bleach, 1 to 2 iterations of 300 ms each) using a LSM Leica SP8 equipped with a 63X N.A. 1.4 PlanApo objective. Confocal stacks were acquired every 2 min after photoconversion and imaged at 22-25°C.

## Quantification and statistical analysis

### Midbody tracking

The apico-basal position of the midbody was calculated measuring the distance between the middle of the new Adherens junction and the midbody (both labeled with MyoII∷RFP) at each time. The x, y, z coordinates of AJ and the midbody were manually tracked to record positions at each time, and then were used to calculate the distance using the Pythagorean Theorem. The midbody apico-basal tracking movement was calculated using the equation y=− ax + b, where a corresponds to the midbody velocity toward the basal pole.

### Signal recovery upon photobleaching

For each FRAP experiment, three mesures were performed, the photobleached junction, the control junction (the finger-like protrusion opposite to the FRAPed finger-like protrusion) and the background.

Data were normalized using EasyFRAP-web software with the “Full scale” method. Signal recoveries were approximated with the equation y= ymax (1-ekxt) with the half-life t1/2 = ln (2) / k.

### Statistical tests

All information concerning the statistical tests are provided in the main text and in the figure legends, including the number of samples analyzed in each experiment. Prism 8 software or R 4.2.1 were used to perform the analyses.

Line plots use the following standards: thick lines indicate the means. Bar plots represent mean and errors bars represent the SD.

Shapiro-Wilk normality test was used to confirm the normality of the data and the F-test to verify the equality of SD. The statistical differences of Gaussian data sets were analyzed using the Student unpaired two-tailed t test.

For the analysis of the midbody displacement, we performed an ANCOVA to test the effect of the interaction between time (between 5 and 120 min) and the « condition » parameter (SOP vs EC). For the midbody displacement and FRAP, the colored areas show standard deviation (SD).

Time to cytoplasmic isolation was modeled via logistic regression, using the GLM function in R, with a binomial error distribution and a logistic link function. We began with a fully parameterized model in which cytoplasmic isolation varied as a function of time, condition and the interaction between time and condition (i.e. both the slopes and intercepts of the model describing temporal effects differed among condition groups). Statistical significance of each model term was evaluated via analysis of deviance, with forward model selection of terms and goodness-of-fit assessed against a Chi-squared distribution. Additionally, we ran two additional nested models: one excluding the interaction term (i.e. a common model slope with differences amongst condition groups in their intercept terms; equivalent to differences in the means amongst groups), and a second in which cytoplasmic isolation state varied only as a function of time (i.e. no effective differences amongst condition groups). We compared overall fit for each model using Akaike’s information criterion (AIC), selecting the parsimony model as the one with the lowest AIC score. Results of analysis of deviance and model comparisons corroborated each other, and so the parsimony model was used to predict the mean percentage of cells demonstrating cytoplasmic isolation, as well as standard errors (SE) of the estimates, and to visualize differences among groups. In some lines, the interaction between time and treatment was not significant, however the interaction between sca> and pnr> was slightly significant (Table S1). These results were corroborated by model comparison via AIC, with the selection process favoring models including only time and treatment effects, i.e. although proportions of cells in cytoplasmic isolation differed among conditions at a given time point, the rate of relative increase over time did not differ among groups. This lack of significant time-treatment interaction facilitated the comparison of treatment effect size. To do so, we used model results to estimate the time at which 50% of the pIIa/pIIb cell indicated cytoplasmic isolation (herein termed t1/2). Each t1/2 were estimated using the two models (sca> and sca>shrubRNAi) and (pnr>, pnr>+ ShrubGFP/+, pnr>+ Cora RNAi, pnr>Cora RNAi+ ShrubGFP/+). Statistical significance were represented as follow: p-value>0.05 ns (not significant); p-value≤0.05*; p-value≤0.01** and p-value≤0.001***

---

**Fig. S1: Midbody assembly and basal displacement throughout EC cytokinesis**

(A) Time-lapse of EC dividing cells (n=20) expressing MyoIIRFP (magenta), ATPαGFP (green). A’ correspond to the orthogonal views along the white dashed lines depicted at t=10 min on panels A. White arrows point to the midbody. AJ, adherens junction; SJ, septate junction; Ap: Apical; Ba: Basal. Time is in minutes with t=0 corresponding to anaphase onset and scale bars represents 5 μm.

**Fig. S2: Evolution of SJ-positive finger-like protrusions and midbody contacts throughout EC cytokinesis**

(A) Time lapse of EC (A, n=13) dividing cells expressing MyoIIRFP (magenta), nlsRFP (magenta) close to cells expressing MyoIIRFP (magenta) and ATPαGFP (green, grey bottom panels). White dashed lines delineate the clone border at t0. White arrows point to the midbody. AJ, adherens junction; SJ, septate junction. Time is in minutes with t0 corresponding to the onset of anaphase. Scale bar represents 5 μm.

**Fig. S3: Maturation of the midbody and timing of cytoplasmic isolation**

(A) Schematic representation of KAEDE photoconversion in SOP cells. The two daughter cells (pIIa and pIIb) are both expressing KAEDE (Green). After photoconversion in pIIa cell, Green KAEDE is photoconverted into Red KAEDE (Magenta). 2 min to until its equilibrium at 8 min after photoconversion, Red KAEDE diffuses in the pIIb cell (No cytoplasmic isolation) or do not diffuse into pIIb (Cytoplasmic isolation). adapted from (Houssin et al., 2021)

(B) Frequency of pIIa-pIIb control cells *pnr*> (black) and *sca*> (grey) photo-converted at different time after anaphase onset.

(C) Plot representing the proportion of pIIa-pIIb cells to have cytoplasmic isolation over time after anaphase onset in control *pnr*> (*pnr*>, dashed line, n=62, 24 pupae) and control *sca*> (*sca*>, solid line, n=30, 11 pupae). Cytoplasmic isolation was assessed based on the ability of photo-converted KAEDE in pIIa/pIIb cell to diffuse in the pIIb/pIIa cell at different time point after the onset of anaphase. Lines represent mean values, predicted from a GLM with the interaction between time and conditions; SE of the estimates are represented in grey shading.

(D) Frequency of pIIa-pIIb control cells (*sca*>) photo-converted at different time after anaphase onset.

(E) Time lapse of SOP dividing cells expressing MyoIIRFP (magenta), MyoIIGFP (green) and GAP43IR (grey). White arrows point to the midbody. (E’) Higher magnification of the white dashed square depicted at t0 in (E) shows the midbody.

(F) Time lapse of SOP dividing cells expressing MyoIIRFP (magenta), PavGFP (green) and GAP43IR (grey). White arrows point to the midbody. (F’) Higher magnification of the white dashed square depicted at t0 in (F) shows the midbody. Time is in minutes. Blue dots mark the pIIa (E, F). Scale bars represent 5 μm (E-F’) and 1 μm (E’, F’).

**Fig. S4: Recruitment of Shrub at the midbody and function of Shrub in cytoplasmic isolation**

(A) Time lapse of EC dividing cell expressing MyoIImcherry adjacent to cells expressing MyoIImcherry (magenta) together with ShrubGFP expressed under *Ay* Gal4 driver (green). White line delineates the border of clones of cells expressing ShrubGFP at t0. The higher magnifications at the bottom right (MB level, t10-36) correspond to the ROI in the white dotted line square of the corresponding panel (scale bar represents 1 μm). (A’) Representation of the EC at t36 (A) surrounded by EC positive for ShrubGFP (light green) and showing the ShrubGFP punctae (dark green) recruitment at the tip of the finger-like protrusion pointing toward the midbody (magenta square).

(B) Time lapse of dividing EC expressing MyoIImcherry (magenta) together with ShrubGFP under the *Ay* Gal4 driver (Green) close to cells expressing MyoIIRFP (magenta) but not ShrubGFP. White dashed line delineates the clone border at t=0. The higher magnifications at the bottom right (MB level, t12-90) correspond to the ROI in the white dotted line square of the corresponding panel (scale bar represents 1 μm). (B’) Representation of the ShrubGFP positive EC (light green) at t60 (B) surrounded by ECs negative for ShrubGFP and showing the ShrubGFP puncta (dark green) recruitment at the midbody (magenta square).

(C) Localization of KAEDE (green) and HRS (magenta) in nota expressing Shrub RNAi and KAEDE. The white dashed lines separate control from Shrub RNAi (KAEDE expressing cells).

(D) Localization of KAEDE (green) and NrxIV (magenta) in nota expressing Shrub RNAi and KAEDE. The white dashed lines separate control from Shrub RNAi (KAEDE expressing cells).

(E) Localization of KAEDE (green) and Sinus (magenta) in nota expressing Shrub RNAi and KAEDE. The white dashed lines separate control from Shrub RNAi (KAEDE expressing cells).

(F) Higher magnification of a SOP midbody (magenta) showing CHMP2BGFP recruitment (green).

AJ, adherens junction; MB, Midbody. Time is in minutes with t0 corresponding to anaphase onset (A-B’). Scale bars represent 5 μm (A, B, C-E) and 1 μm (F).

**Fig. S5:**

(A) Z-projection of confocal images obtained on control (left) or ShrubGFP/+ ovaries (green, middle and right) stained for α-Spectrin (magenta) to visualize the fusome linking the germline stem cell (GSC) to its progeny. In control, the GSC is only linked to its daughter cystoblast (CB). In ShrubGFP/+ ovaries, GSC are either as in control (middle) or linked to several progeny and form stem cysts (right). Scale bars represent 5 μm.

(B) Higher magnification of a SOP midbody (magenta) showing CHMP2BGFP recruitment (green). Scale bars represent 1 μm.

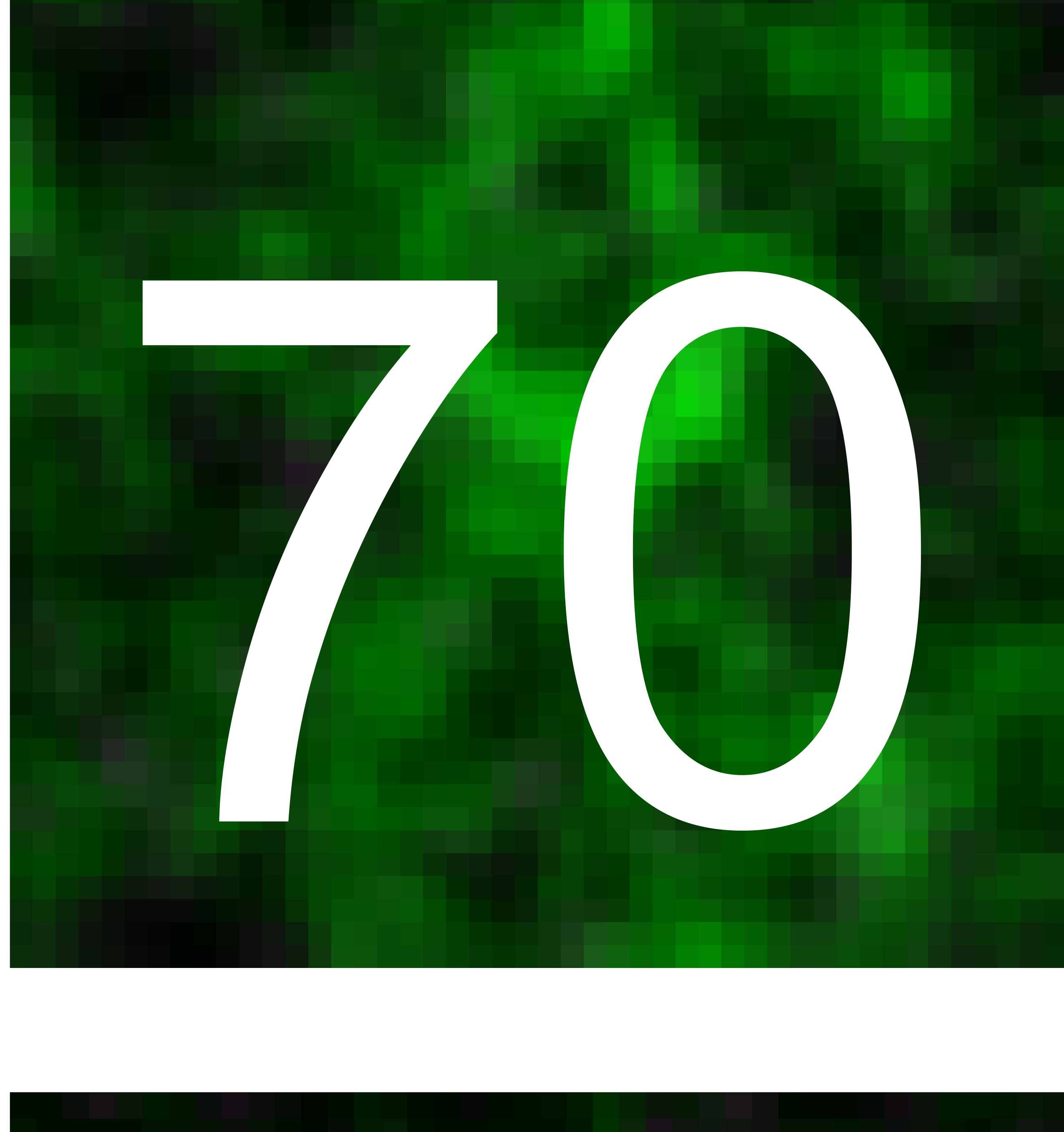

